# Modelling the human IL6RA Asp358Ala variant in mice: effects on atherosclerosis and abdominal aortic aneurysm

**DOI:** 10.64898/2026.10.16.759391

**Authors:** Minoru Takaoka, Rahul Sharma, Fabien Lareyre, Marc Clement, James Harrison, Leanne Masters, Yuning Lu, Dirk Paul, Ziad Mallat

**Author notes:** Correspondence and request for materials should be addressed to Ziad Mallat.

## Abstract

The association of IL6RA Asp358Ala variant with reduced risk of atherosclerotic cardiovascular disease (ASCVD) and abdominal aortic aneurysm (AAA) prompted interest in targeting IL6 signalling in these conditions. A recent report that ziltivekimab, a neutralising anti-IL6 antibody, did not reduce major adverse cardiovascular events in patients with ASCVD and chronic kidney disease, has cast doubt on the causal role of IL6 signalling in ASCVD. Here, we show that introducing the orthologous IL6RA Glu357Ala substitution in male mice leads to the same alteration in soluble IL6RA levels seen in humans, and significantly prevents atherosclerosis and AAA. Thus, therapies that more closely reproduce the biological effects of the IL6RA Asp358Ala variant on IL6 signalling are needed to reduce cardiovascular risk in humans.

## Introduction

Atherosclerosis is a lipid-driven inflammatory disease of the artery wall. Biomarkers of systemic inflammation are associated with cardiovascular outcomes. Two of the most validated inflammatory biomarkers are hsCRP and IL6, with hsCRP being downstream of IL6 membrane receptor (classic) signalling^1^. Mendelian randomization studies have shown that hsCRP is unlikely to be causally involved in ASCVD. However, an IL6RA Asp358Ala genetic variant that results in the shedding of membrane IL6RA, and consequently reduces IL6 classic signalling and limits CRP production, is causally associated with reduced risk of ASCVD^2^. The identification of IL6 signalling as a potential therapeutic target strongly supported the development of IL6 inhibitors for the treatment of ASCVD. The recent announcement that ziltivekimab did not reduce cardiovascular outcomes in patients with ASCVD, chronic kidney disease, and increased hsCRP appeared at odds with the available genetic evidence. Given that the genetic evidence supporting causality may still be influenced by unrecognised pleiotropy, a key question is whether there is unequivocal evidence that the IL6RA genetic variant truly alters ASCVD. Currently, there is no evidence that directly manipulating the genome to introduce this specific genetic variation in IL6RA would directly alter atherosclerosis development. To this end, we have modelled the human IL6RA Asp358Ala genetic variant by generating a knock-in mouse line (IL6RA E357A) carrying the orthologous Glu357Ala substitution. The mice were crossed to low-density lipoprotein receptor deficient (*Ldlr*^*-/-*^) mice to assess the impact of the IL6RA variant on atherosclerosis. The same genetic variant is associated with reduced risk of AAA in humans^3^. Therefore, we also tested the knock-in mouse line in a validated mouse model of AAA.

## Results

The IL6RA E357A mouse line was generated by the Wellcome Sanger Institute, UK (see Online Methods). Genotyped wild-type (Glu/Glu), heterozygous (Glu/Ala), and homozygous (Ala/Ala) littermate mice were used for phenotyping. Similar to the reported increase of soluble IL6RA associated with the Asp358Ala allele in humans^2^, mice expressing the orthologous Glu357 Ala allele showed an allele-dependent increase in circulating soluble IL6RA, without any significant change in soluble gp130 (Figure 1). Challenging the mice with IL1β led to a slightly higher peak of circulating IL6 in Ala/Ala mice (Figure 1), also in agreement with the human data^2^. Therefore, the increase of soluble IL6RA associated with the Asp358 Ala allele in humans is a direct result of the single-point mutation in IL6RA.

**Figure 1.**
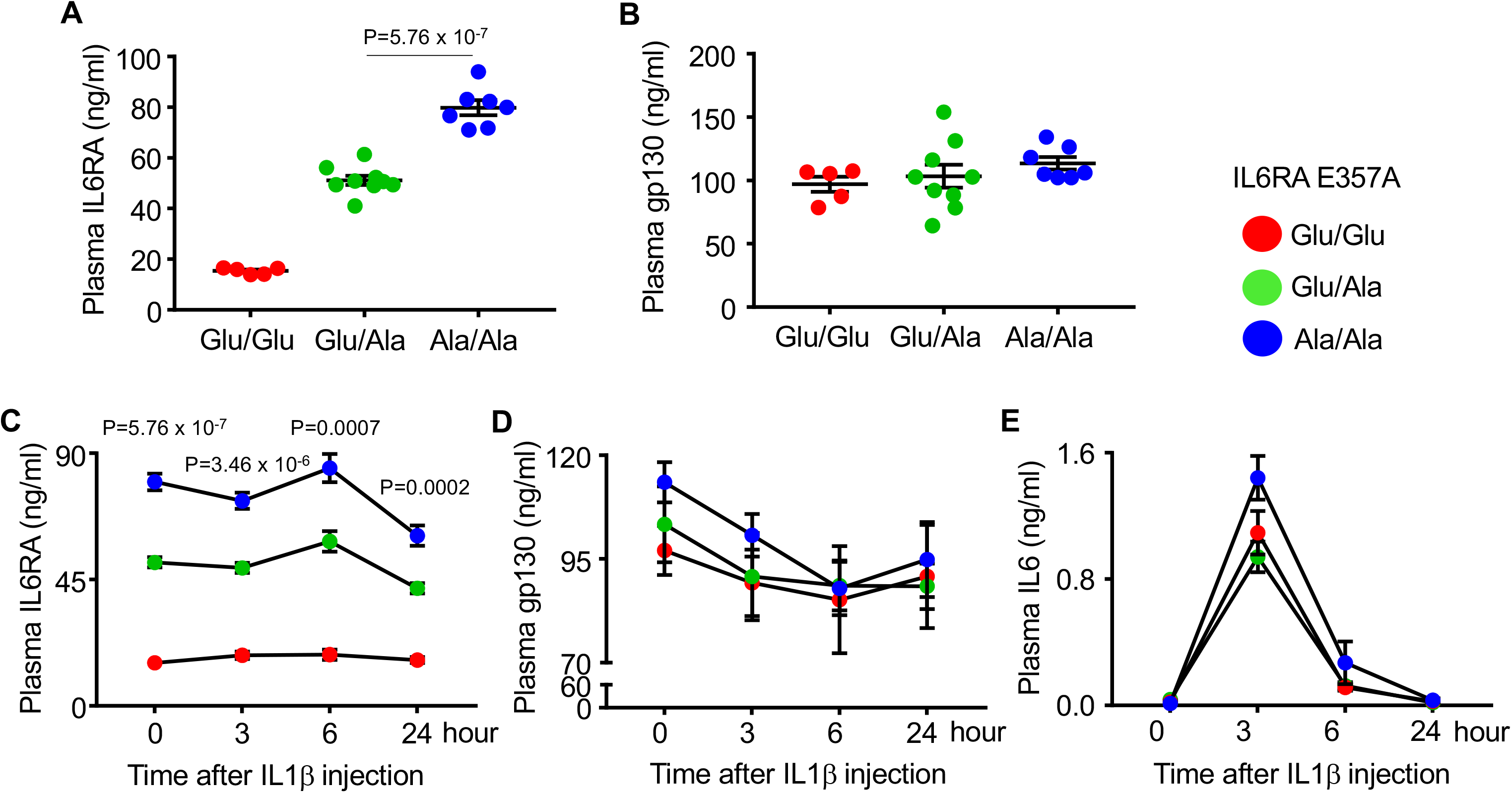
Circulating levels of components of the IL6R signalling pathway in IL6RA E357A knock-in mice. A. Baseline plasma levels of soluble IL6RA (n=5 Glu/Glu, n=9 Glu/Ala, n=7 Ala/Ala genotypes). B. Baseline plasma levels of soluble gp130. C-E. Plasma levels of soluble IL6RA (C), soluble gp130 (D), and IL6 (E) at baseline and after administration of IL1β. Wild-type (Glu/Glu), heterozygous (Glu/Ala), and homozygous (Ala/Ala) littermate mice were used for phenotyping. For A-E, n=5 Glu/Glu, n=9 Glu/Ala, n=7 Ala/Ala genotypes.

To address the impact of the IL6RA variant on atherosclerosis, IL6RA E357A mice were crossed with *Ldlr*^*-/-*^ mice. Atherosclerosis extent in the aortic sinus was assessed in Glu/Glu, Glu/Ala, and Ala/Ala littermate *Ldlr*^*-/-*^ mice after 10 weeks on Western diet. The IL6RA variant did not alter plasma total cholesterol levels in either males (27.28±5.87 mmol/L in Glu/Glu; 24.20±3.79 in Glu/Ala; 23.24±5.18 in Ala/Ala) or females (19.49±1.99 mmol/L in Glu/Glu; 17.05±2.25 in Glu/Ala; 19.58±2.38 in Ala/Ala). However, Ala/Ala *Ldlr*^*-/-*^ male mice showed a significant reduction of atherosclerosis compared with both WT Glu/Glu and heterozygous Glu/Ala genotypes (Figure 2). Lesion size in Glu/Ala *Ldlr*^*-/-*^ male mice was similar to that of Glu/Glu *Ldlr*^*-/-*^ male mice (Figure 2). The IL6RA variant did not alter the development of atherosclerosis in *Ldlr*^*-/-*^ female mice (Figure 2).

**Figure 2.**
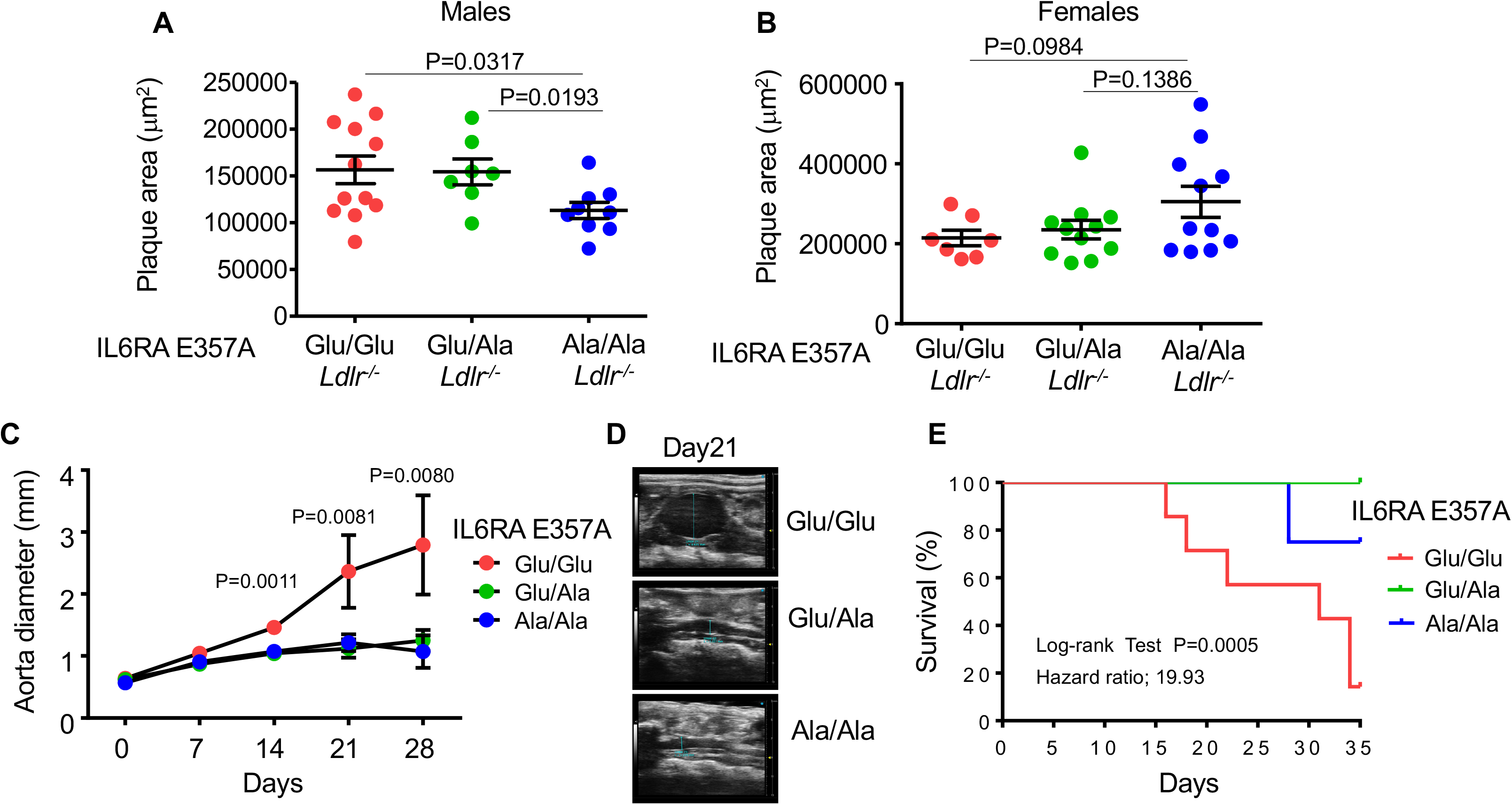
Impact of IL6RA E357A substitution on the development of atherosclerosis and abdominal aortic aneurysm in mice. Atherosclerotic plaque size in the aortic sinus of male (A) and female (B) *Ldlr*^*-/-*^ mice harbouring the wild-type Glu/Glu (n=12 males and 7 females), the heterozygous Glu/Ala (n=7 males and 11 females), or the homozygous Ala/Ala (n=9 males and 11 females) IL6RA genotype. Mice were fed a Western diet for 10 weeks. C-E. Male C57Bl/6N littermate mice harbouring the wild-type Glu/Glu (n=7 Glu/Glu), the heterozygous Glu/Ala (n=8 Glu/Ala), or the homozygous Ala/Ala (n=4 Ala/Ala) IL6RA genotype were subjected to a validated model of severe AAA (see Methods). Abdominal aortic diameter was assessed at baseline and post-AAA induction using serial vascular ultrasound. Quantification of aortic abdominal diameter is shown in (C) and representative pictures of 2D ultrasound imaging of the abdominal aorta are shown in (D). F. Kaplan-Meier curves of survival free from fatal aortic rupture.

Male Glu/Glu, Glu/Ala and Ala/Ala littermate mice were then used in a validated model of severe AAA^4^. As expected, Glu/Glu mice developed severe AAA with substantial abdominal aortic dilatation (Figure 3). In contrast, mice with the Glu/Ala and Ala/Ala genotypes were significantly protected from the development of AAA (Figure 3). Interesting, mice with the Glu/Ala and Ala/Ala genotypes were also substantially protected from the occurrence of fatal aortic rupture (Figure 3).

## Discussion

Despite human genetic evidence of an association of the IL6RA Asp358Ala variant with the risk of ASCVD and AAA, there was still no experimental evidence supporting a direct causal link between the genetic variant and disease development. Such direct experimental evidence remains essential given that pleiotropy can never be excluded in Mendelian randomisation analyses. Here, we show that introducing the human orthologous IL6RA Glu357Ala substitution in mice leads to an Ala allele-dependent increase in soluble IL6RA, as previously described in humans, and significantly reduces the development of atherosclerosis and AAA in validated mouse models. Heterozygous Glu/Ala mice were not protected from atherosclerosis during this relatively short exposure to the Western diet. A longer-term exposure may be needed to unravel a protective effect. Whether the presence of the IL6RA Asp358Ala variant associates with the extent of human atherosclerosis is currently unclear. However, IL6RA Asp358 allele was shown to be associated with a particular carotid atherosclerosis endotype characterised by increased expression of OncoStatinM Receptor, which conferred an increased risk of myocardial infarction compared with individuals harbouring the IL6RA Ala allele^5^.

For AAA, the association with the IL6RA Asp358Ala variant is stronger than for ASCVD, with approximately 10% lower AAA risk per copy of the Ala allele^3^, compared with only 3.4% lower ASCVD risk^2^. This is reflected in our experimental data, with mice bearing one copy of the Ala allele being already protected from AAA. The available human data are not conclusive regarding the association of IL6RA Asp358Ala with the risk of AAA progression^6^ or rupture^3^, mostly due to limited sample size. Our experimental data indicate that the orthologous IL6RA Glu357Ala substitution directly protects mice from the risk of AAA rupture. The association of IL6RA Asp358Ala with the risk of AAA appears to be stronger in men than in women^3^, but no such sex-dependent analyses are currently available for the association with ASCVD. However, our observation that female mice with the IL6RA Ala357 allele are not protected from atherosclerosis suggests a sex-dependent mechanism. Whether this is related to the duration of exposure to the genetic variant, the stage of lesion development, or the influences of sex hormones on the disease process will require further studies.

The human genetic evidence supporting a causal role of IL6 signalling in ASCVD prepared the ground for the development of IL6 inhibitors to tame the persistent inflammatory risk associated with adverse cardiovascular outcomes. Therefore, the neutral result of the ZEUS trial was unexpected. By providing direct experimental evidence that the IL6RA 357/358Ala variant is causally related to protection against ASCVD, our data re-affirm the importance and relevance of targeting IL6 signalling in ASCVD. Importantly, while ziltivekimab blocks IL6 classic and trans-signalling pathways, the only well-documented perturbations of IL6 signalling associated with the IL6RA variant are those related to IL6 classic signalling^7^. The impact of the IL6RA variant on IL6 trans-signalling and IL6 trans-presentation remains unclear and requires further studies. The knock-in IL6RA Glu357Ala mice described here will be extremely valuable in pinpointing the distinct roles of the different modes of IL6 signalling in the context of ASCVD and AAA, as well as in other disease settings influenced by the IL6RA Asp358Ala variant. Using these mice, our results already suggest that therapies that mimic the biological effects of the IL6RA Asp358Ala variant, as could be achieved through targeted shedding of IL6RA by induced protease recruitment^8^, are likely to reduce cardiovascular risk in humans.

In summary, IL6RA Glu357Ala substitution in male mice phenocopies the impact of the human IL6RA Asp358Ala variant on soluble IL6RA levels, and leads to a significant reduction of the development of atherosclerosis and abdominal aortic aneurysm.

## Methods

### Mice

Mice carrying the IL6R E357A substitution (originally named Il6ra^em1(IMPC)Wtsi^ allele) were generated by the Wellcome Sanger Institute, UK, on a C57BL/6N genetic background using CRISPR-mediated genome editing. The allele introduces the E357A substitution in murine IL6RA, corresponding to the human IL6RA Asp358Ala variant (rs2228145). Genotyping was performed using custom TaqMan SNP assay (Thermo Fisher Scientific, Assay ID ANFVP4C, Cat. No.4332077) on a Roche Lightcycler or Applied Biosystems 7500 Fast PCR System. Allelic discrimination and genotype clustering analysis were performed by measuring the relative fluorescence units of the FAM and VIC reporter dyes. The Sanger technical datasheet documents the targeted mutation, guide RNA and donor oligonucleotide, and genotyping assays. The mouse line was available through the European Mouse Mutant Archive (EMMA ID: EM:13825; RRID:IMSR_EM:13825). Il6raE357A mice were crossed with *Ldlr*^*-/-*^ mice to generate Glu/Glu wild-type, Glu/Ala heterozygous, and Ala/Ala homozygous littermate mice on *Ldlr*^*-/-*^ background.

### Atherosclerosis model

The atherosclerosis experiments were conducted on both male and female Glu/Glu wild-type, Glu/Ala heterozygous, and Ala/Ala homozygous littermate *Ldlr*^*-/-*^ mice. Mice aged 6 to 12 weeks were fed a Western diet (21% fat, 0.15% cholesterol) for 10 weeks. At the end of the experiment, arteries and hearts were collected into 4% PFA overnight before washing into PBS. Aortic root cryosections were stained with Oil Red O (Sigma-Aldrich, O0625) for atherosclerotic lesion size. Images were captured and analysed using a Leica DM6000B microscope and ImageJ (NIH).

### Aortic aneurysm and dissection model

Male IL6RA E357A Glu/Glu wild-type, Glu/Ala heterozygous, and Ala/Ala homozygous littermate mice aged 12-14 weeks were used. Elastase (Sigma; Type I, ≥4.0 units/mg protein) was applied topically to the abdominal aorta, and starting on the day of surgery, 250 μg of a mouse anti-mouse TGFβ antibody (BioXcell; clone 1D11.16.8) was administered intraperitoneally 3 times a week for 5 weeks, as previously described (28912363).

### Aortic ultrasound

Aortic diameters were measured in mice anesthetized with 2% isoflurane using the Vevo 3100 ultrasound system (VisualSonics; 30 MHz MX400 probe). To evaluate the maximum cross-sectional diameter, B-mode sagittal images were acquired at the site of maximal dilation of the AAA.

### Biomarker measurements

Plasma IL6RA, gp130, and IL6 concentrations were measured using mouse ELISA kits (Thermo Fisher, EMIL6RA, EMIL6ST, and BMS603-2) in the presence or absence of stimulation with recombinant mouse recombinant mouse IL-1β protein (Biolegend, 575104).

### Statistical Analysis

For the atherosclerosis experiments, sample size calculation was based on the requirement to detect 30% difference in lesion size between WT Glu/Glu and homozygous Ala/Ala *Ldlr*^*-/-*^ mice. Sample size calculation did not take into account the heterozygous Glu/Ala *Ldlr*^*-/-*^ mice because we expected atherosclerotic disease to be less affected in the heterozygous mice compared with the homozygous mice during a relatively short duration (10 weeks) of Western diet. In humans, a meta-analysis reported only a 3.4% lower ASCVD risk per copy of the Ala allele over a lifetime of exposure (22421339). Therefore, the main comparison was between Glu/Glu and Ala/Ala *Ldlr*^*-/-*^ mice. However, we pre-specified secondary comparisons between Glu/Glu and Glu/Ala *Ldlr*^*-/-*^ mice and between Glu/Ala and Ala/Ala *Ldlr*^*-/-*^ mice. For an expected mean lesion size of 175,000 mm^3^ and SD of 40,000, a sample size of n=10 per group was needed to detect 30% difference in lesion size with 80% power and two-sided type I error of 0.05. The AAA experiment was designed as an exploratory preliminary study. Therefore, no sample size calculation was conducted a priori. Available Glu/Glu, Glu/Ala and Ala/Ala littermate male mice were used for the experiment.

## Acknowledgements

This study was supported by the British Heart Foundation (/10/001/27642), the Leducq Foundation (ERAADICATE) and the BHF Cambridge Centre of Excellence (RE/24/130011).

## Author contributions

MT performed the atherosclerosis experiments. RS and FL performed the AAA and ultrasound experiments. MC, JH, LM and YL helped with both the atherosclerosis and AAA experiments and performed genotyping and biomarker measurements. DP was involved in the design of the knock-in mouse model. ZM conceptualised and supervised the work, wrote the manuscript and provided funding. All co-authors reviewed the manuscript for accuracy and intellectual 202 content.

